# Is the neuropeptide PEN a ligand of GPR83?

**DOI:** 10.1101/2023.08.31.555736

**Authors:** Yvonne Giesecke, Vahid Asimi, Valentina Stulberg, Gunnar Kleinau, Patrick Scheerer, Beate Koksch, Carsten Grötzinger

**Affiliations:** Tumor Targeting Group, Department of Hepatology and Gastroenterology, Charité - Universitätsmedizin Berlin, 13353 Berlin, Germany; Institute of Chemistry and Biochemistry, Freie Universität Berlin, 14195 Berlin, Germany; Group Structural Biology of Cellular Signaling, Institute of Medical Physics and Biophysics, Charité – Universitätsmedizin Berlin, 10117 Berlin, Germany

**Keywords:** GPR83, PEN, proSAAS, orphan receptor, GPCR, peptide

## Abstract

G protein-coupled receptor 83 (GPR83) is a class A G protein-coupled receptor with predominant expression in the cerebellum and proposed function in the regulation of food intake and in anxiety-like behavior. The neuropeptide PEN has been suggested as a specific GPR83 ligand. However, conflicting reports exist about whether PEN is indeed able to bind and activate GPR83. This study was initiated to evaluate PEN as a potential ligand of GPR83. Employing several second-messenger and other GPCR activation assays as well as a radioligand binding assay, and using multiple GPR83 plasmids and PEN peptides from different sources, no experimental evidence was found to support a role of PEN as a GPR83 ligand.

## 1. Introduction

G protein-coupled receptor 83 (GPR83), also termed glucocorticoid-induced receptor (GIR), GPR72, KIAA1540, or JP05, is a class A G protein-coupled receptor with predominant expression in the cerebellum and proposed function in the regulation of food intake and in anxiety-like behavior [1-3]. A metabolic role was supported by studies demonstrating significant expression in brain areas associated with food intake [2,4,5]. A potential role of GPR83 in regulatory T cell development and function has not been confirmed [6-8]. Recently, GPR83-expressing projection neurons were shown to be involved in conveying thermal, tactile and noxious cutaneous signals from the spinal cord to the brain [9]. The identity of a ligand for GPR83 remained unknown until recently. In 2016, it was proposed that the neuropeptide PEN, which is released from the proprotein PCSK1N or proSAAS through posttranslational processing, serves as a distinct ligand for GPR83 [10]. A previous report has implicated PEN in the regulation of feeding behavior [11]. Independent reports from two other labs also supported activation of GPR83 by PEN [12,13]. In addition, a previously unreported 9-kDa protein termed family with sequence similarity 237 member A (FAM237A) was revealed to activate GPR83 in a peptide hormone library screening study [14]. Finally, a procholecystokinin (proCCK)-derived peptide, proCCK56-63 with sequence similarity to PEN, was recently introduced as a ligand and agonist of GPR83 [15].

A 2023 paper, however, has comparatively evaluated the capacity of all three ligands proposed so far (PEN, proCCK56-63 and FAM237A) to bind and activate GPR83. Using multiple assay systems, only FAM237A was confirmed as a GPR83 ligand and agonist, but not PEN or proCCK56-63 [16]. The present study was initiated to evaluate PEN as a ligand of GPR83. Employing several second-messenger and other GPCR activation assays as well as a radioligand binding assay, and using multiple GPR83 plasmids and PEN peptides from different sources, no experimental evidence was found to support a role of PEN as a GPR83 ligand.

## 2. Results

### 2.1. Overexpression of GPR83 and the deletion variant GPR83del in human cell lines

To test for cell-based GPR83 activation and binding, cell models were created by transfecting cells and selecting overexpressing clones of the human embryonic kidney cell line HEK293 and the human osteosarcoma cell line U2OS. In addition to wild-type GPR83, a deletion variant GPR83del was transfected, which almost completely lacks the aminoterminal ectodomain, found to show increased signaling [17]. To monitor expression and localization, both proteins were also expressed with an aminoterminal hemagglutinin peptide tag (HA tag). According to mRNA quantification by reverse-transcription quantitative PCR (RT-qPCR), transfection increased GPR83 expression in HEK293 cells approximately 100fold, while the increase in U2OS cells was approximately 1000fold (Figure 1A). Overexpression was further confirmed by detection of HA-tagged GPR83 protein in the appropriate cell clones by immunofluorescence microscopy (Figure 1B).

**Figure 1.**
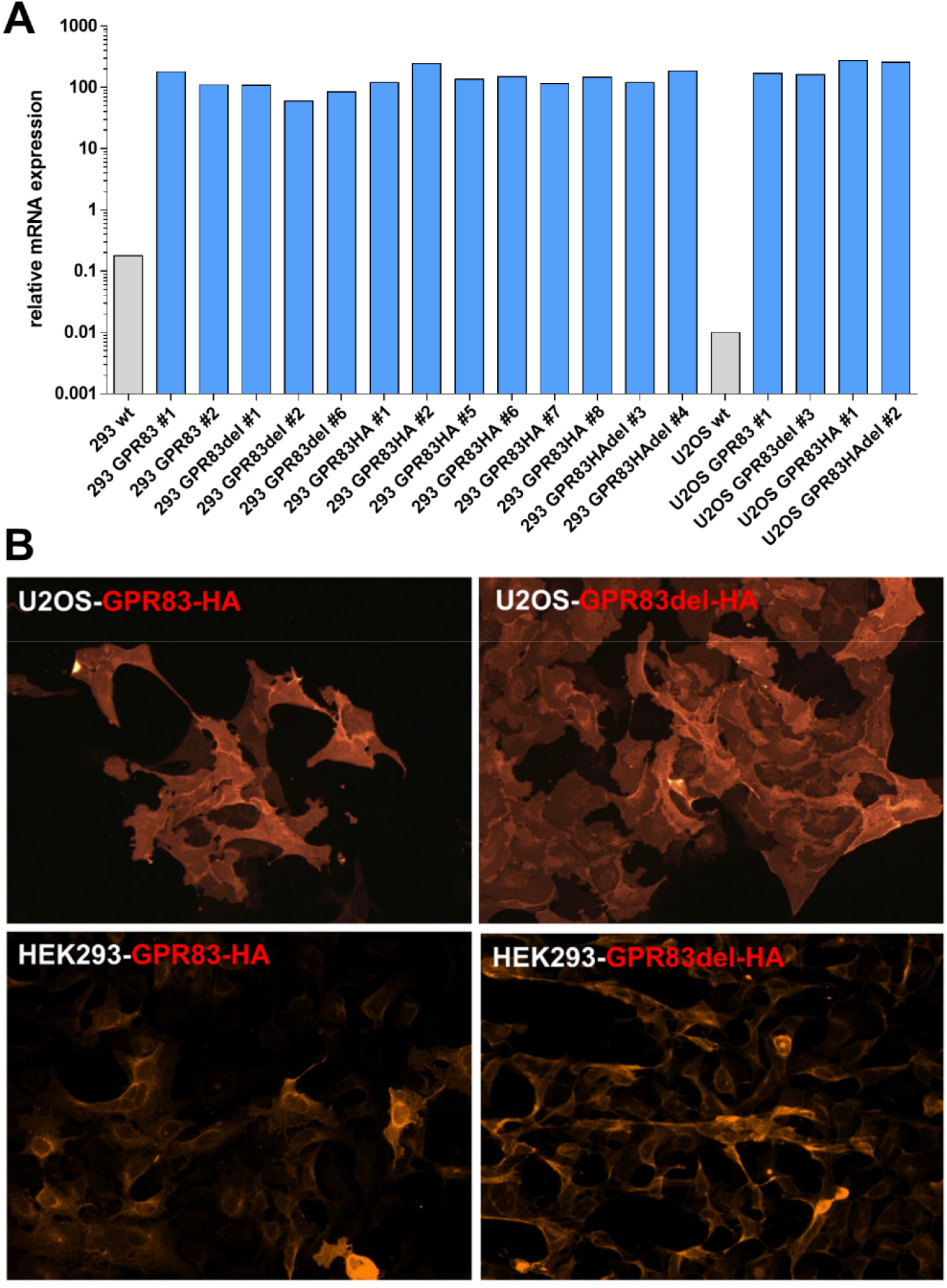
Evidence of overexpression of untagged and HA-tagged GPR83 and the deletion variant GPR83del in two human cells lines. (**A**) Expression as determined by RT-qPCR of GPR83 mRNA in wild-type (wt) HEK293 (293 wt) and U2OS (U2OS wt) cells is shown by grey bars, expression of cell clones transfected with plasmids encoding GPR83, the deletion variant GPR83del and the HA-tagged versions of these (blue bars, designation and clone number stated below). (**B**) Detection of overexpressed HA-tagged GPR83 and GPR83del protein in transfected HEK293 and U2OS cell clones by immunofluorescence microscopy.

### 2.2. Intracellular calcium mobilization

As activation of many GPCRs is coupled to calcium mobilization from intracellular stores, GPR83 was probed for ligand-dependent calcium release. Wild-type HEK293 cells and HEK293 clones overexpressing either full-length GPR83 or the N-terminal deletion variant GPR83del were exposed to buffer or one of two independent synthesis batches (PEN #1, PEN #2) of the neuropeptide PEN, while calcium indicator fluorescence was recorded. While the positive control substances (co mix [carbachol, ATP, dimaprit] and ionomycin) yielded a clear calcium signal above buffer background, PEN did not (Figure 2A). Calcium release was also tested in a second cell line, osteosarcoma U2OS cells, using various stably expressing clones. When added 10 seconds after the start of the recording, positive controls led to a moderate or strong and sustained enhancement of calcium indicator fluorescence, while PEN did not (Figure 2B). The G protein G_α16_ has been used couple GPCRs to the calcium pathway, which do not spontaneously signal via the G_q_ pathway. Therefore, HEK293 cells transfected with G_α16_ were employed to investigate a potential GPR83-dependent calcium mobilization by PEN. The positive controls ATP and carbachol strongly enhanced the calcium signal, whereas PEN did not (Figure 2C). To demonstrate that G_α16_ transfection was functional, two other GPCRs and their cognate ligands were tested and yielded a robust calcium response (Figure 2C). A number of GPCRs show ligand-independent basal calcium activity. However, in an experiment comparing basal calcium levels in wild-type HEK293 cells as well as in HEK293 clones overexpressing four different variants of GPR83, they were not found to be different (Figure 2D).

**Figure 2.**
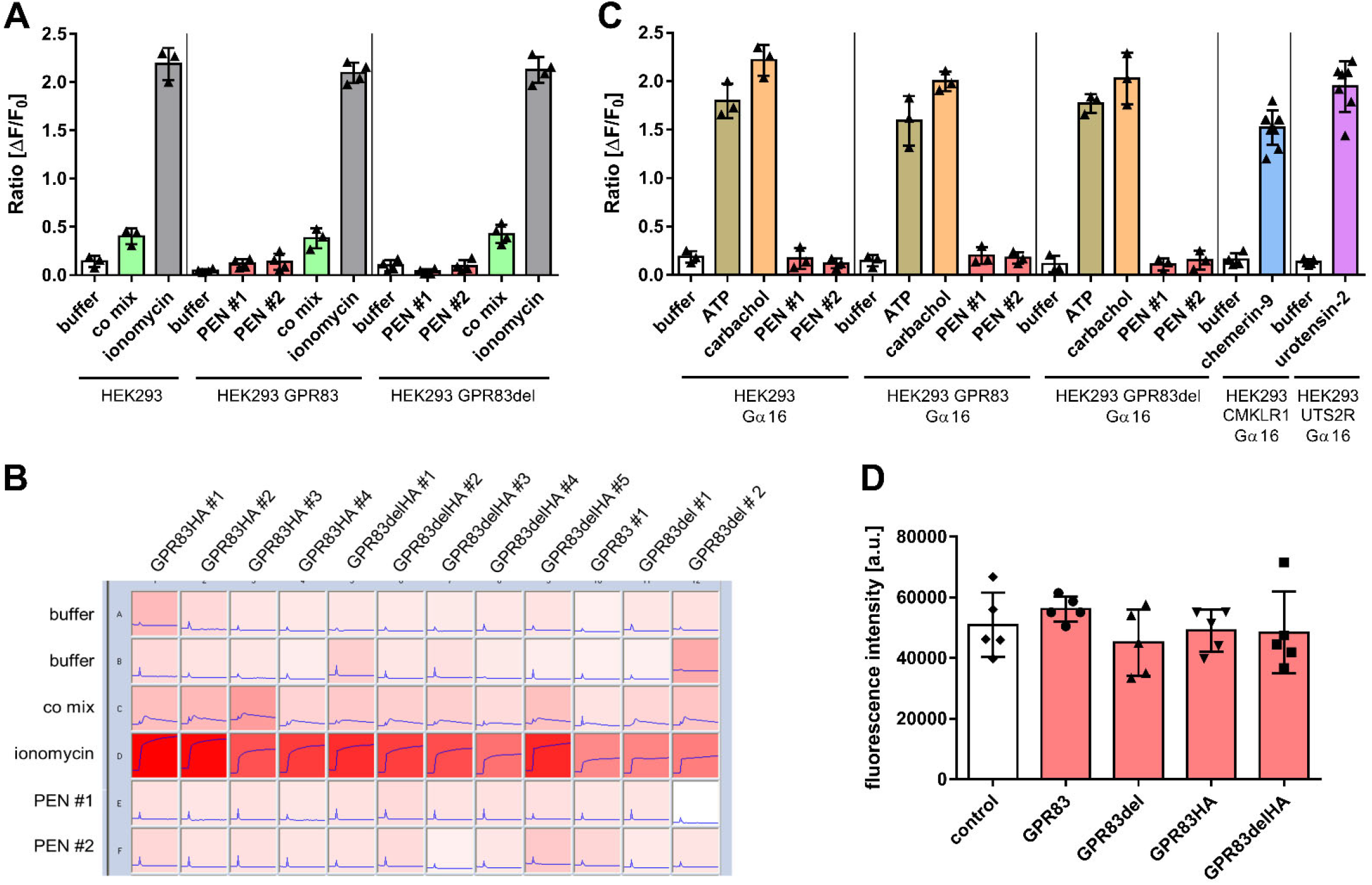
Investigation of ligand-dependent and ligand-independent intracellular calcium mobilization. (**A**) Wild-type HEK293 cells and HEK293 cell clones overexpressing either GPR83 or the deletion variant GPR83del were incubated with either buffer or the specified ligand (10 μM PEN or co mix = 100 μM carbachol, 100 μM ATP, 40 μM dimaprit, or 50 μM ionomycin). **(B)** Calcium indicator fluorescence intensity traces over 60 seconds of various U2OS cell clones overexpressing untagged or HA-tagged GPR83 or the deletion variant GPR83del before and after addition of buffer or ligand (co mix as in (A) or 50 μM ionomycin or 10 μM PEN). **(C)** HEK293 cells overexpressing either G_α16_, GPR83 and G_α16_, the deletion variant GPR83del and Gα16, CMKLR1 and G_α16_, or UTS2R and G_α16_ were incubated with either buffer or the specified ligand (10 μM PEN or 100 μM carbachol or 100 μM ATP). **(D)** Basal calcium indicator fluorescence of wild-type HEK293 cells and HEK293 cell clones overexpressing one of the specified GPR83 proteins.

### 2.3. Ligand-dependent cAMP formation

Production of cyclic adenosine monophosphate (cAMP) is a hallmark of GPCRs signaling via the Gs-induced pathway. To investigate the potential of PEN to stimulate cAMP production mediated by GPR83, levels of the second messenger were determined after incubation with PEN of various GPR83-overexpressing clones as well as wild-type HEK293 and U2OS cells. While the adenylyl cyclase activator forskolin strongly increased cAMP levels (observed as a reduction of the fluorescent signal in the competition assay), PEN did not change it (Figure 3A). Likewise, a titration of ligand over a concentration range from 50 pM to 1 μM in selected GPR83-expressing HEK293 and U2OS clones did not indicate PEN-dependent cAMP formation (Figure 3BC).

**Figure 3.**
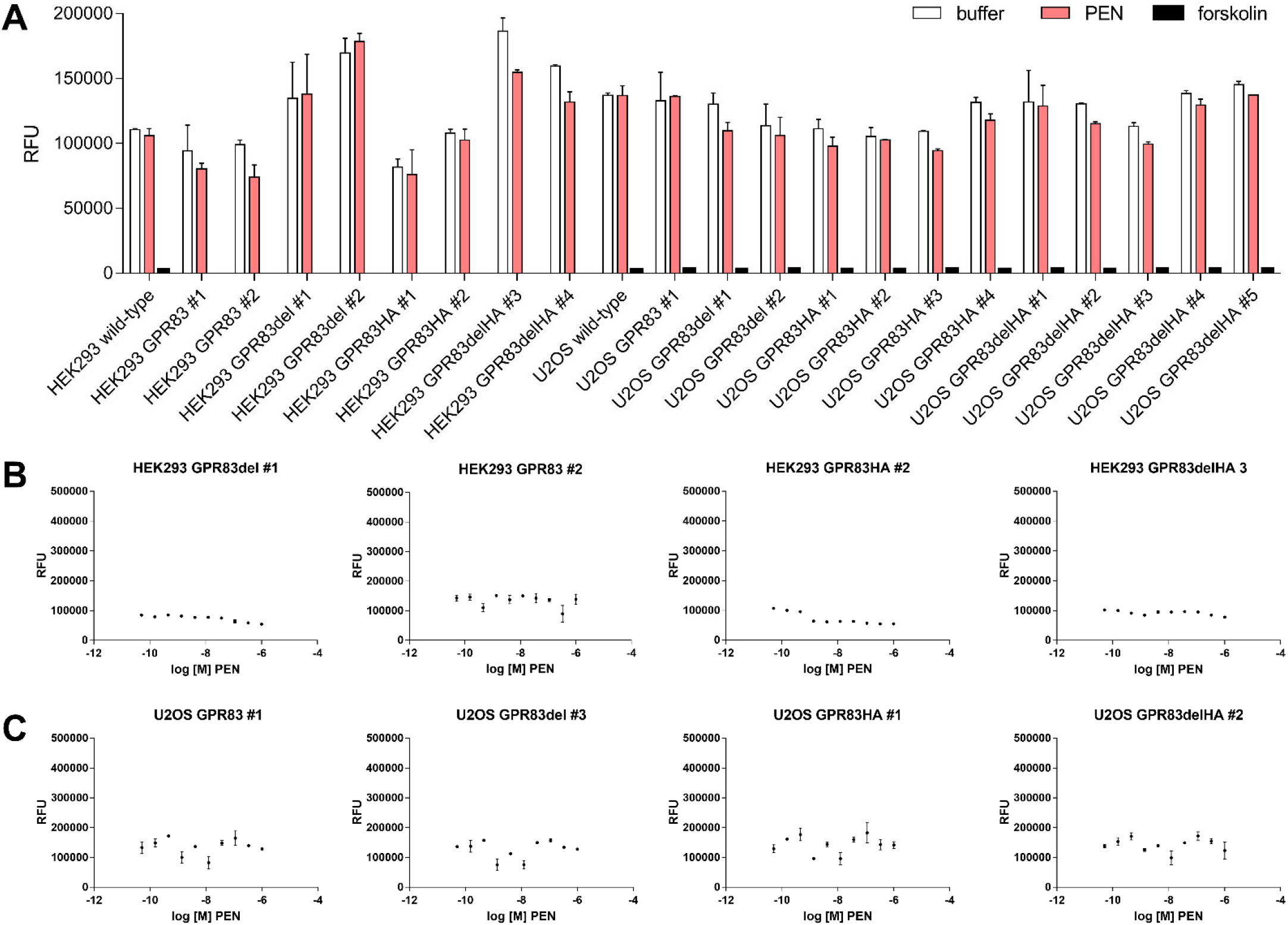
Analysis of ligand-dependent, GPR83-mediated cAMP production. **(A)** Wild-type HEK293 and U2OS cells as well as various clones with confirmed overexpression of either GPR83 or the deletion variant GPR83del (both ± HA tag) were incubated with either buffer, 1 μM PEN or 50 μM forskolin. **(B)** Selected GPR83-overexpressing HEK293 clones were assayed for cAMP production using different concentrations of PEN. **(C)** Selected GPR83-overexpressing U2OS clones were assayed for cAMP production using different concentrations of PEN.

### 2.4. ERK1/2-phosphorylation and β-arrestin activation

Apart from G protein-mediated second messenger signaling via Ca^2+^ and cAMP, GPCR stimulation may also lead to activation of β-arrestin and phosphorylation of extracellular signal-regulated kinases 1/2 (ERK1/2). Immunoblot analysis was employed to investigate ligand-dependent phosphorylation of ERK1/2 in HEK293 cells overexpressing GPR83 or the deletion variant GPR83del as well as in wild-type HEK293 cells. PEN and the positive control substances epithelial growth factor (EGF) and phorbol-12-myristate-13-acetate (PMA) all substantially induced phosphorylation of ERK1/2, with EGF yielding the strongest response (Figure 4A-B). Remarkably, PEN induced an equally strong ERK1/2 phosphorylation in wild-type HEK293 cells as well as in HEK293 cells overexpressing GPR83 or GPR83del, thus independent of GPR83 levels in the cells (Figure 4A-B).

**Figure 4.**
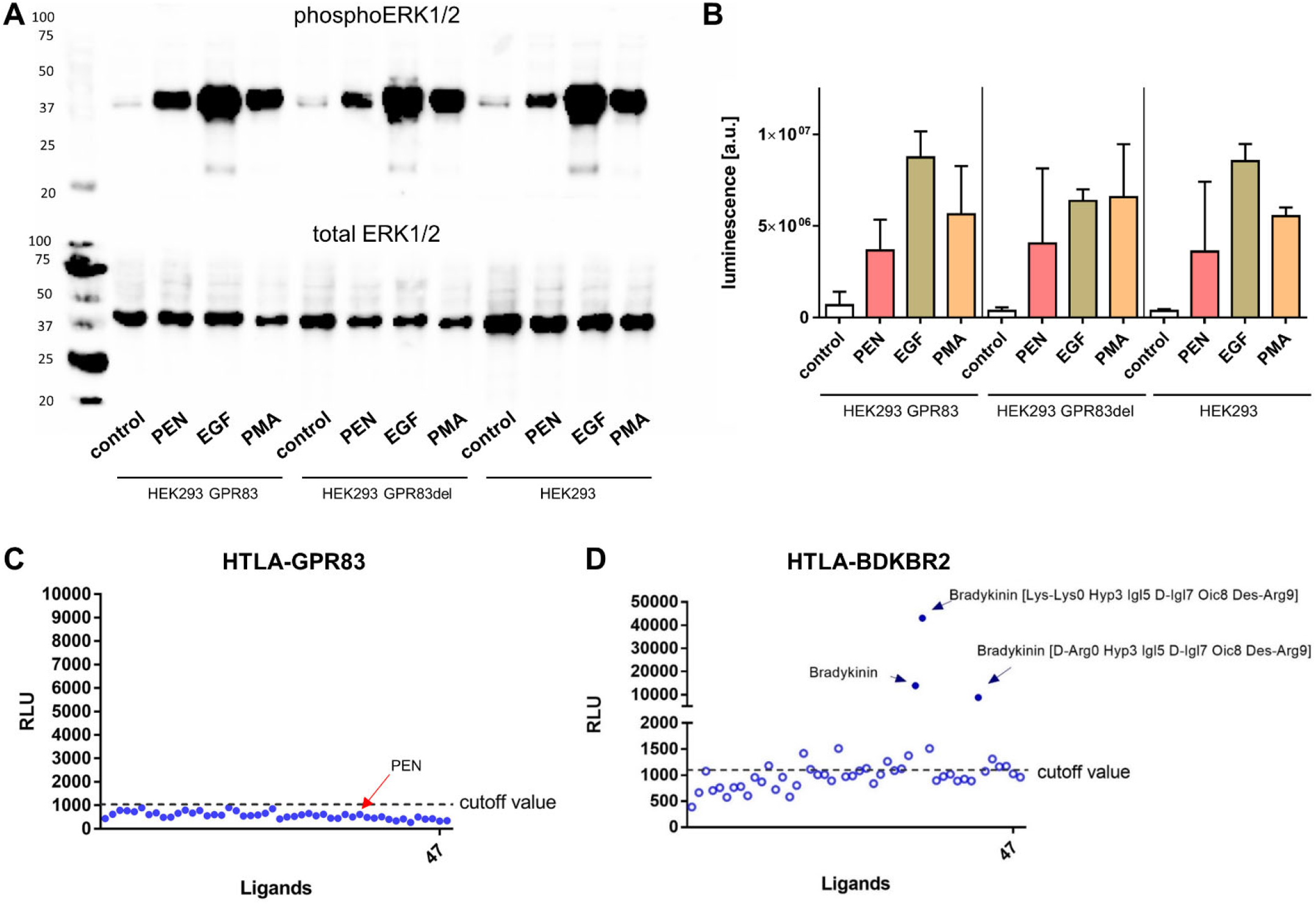
Ligand-dependent phosphorylation of ERK1/2 and β-arrestin (TANGO) assay. **(A)** Immunoblot detection of phosphorylated (upper blot) and total (lower blot) ERK1/2 in lysates of HEK293 wild-type cells and in HEK293 cells overexpressing GPR83 or GPR83del. **(B)** Quantification (mean ± SD of n=2 independent experiments) of chemiluminescent readout of the immunoblots shown in (A). **(C)** TANGO assay for ligand-induced β-arrestin activity in HTLA cells overexpressing GPR83 (relative light units, RLU). **(D)** TANGO assay for ligand-induced β-arrestin activity in HTLA cells overexpressing bradykinin receptor BDKBR2.

The TANGO assay reports GPCR-related activation of β-arrestin via a luminescent readout [18]. When reporter HTLA cells expressing GPR83 were incubated with a ligand library containing PEN, no reactivity was detected (Figure 4C), while HTLA cells expressing bradykinin receptor BDKBR2 yielded strong luminescent responses upon incubation with either of three bradykinin-related ligands contained in the library (Figure 4D).

### 2.5. Radioligand binding

One of the most proven methods for quantitatively analyzing the interaction of biomolecules is the competitive radioligand binding assay. As natural PEN does not contain a tyrosine (Tyr) residue for radiolabeling with ^125^I, it had to be artificially introduced. To safeguard against possible interference of the added Tyr with ligand-receptor binding, two peptides were created: one with the Tyr on the aminoterminal end (Tyr-PEN), and the other with Tyr added on the carboxyterminus (PEN-Tyr). Radiolabeled ^125^I-PEN-Tyr and ^125^I-Tyr-PEN were then used on wild-type HEK293 cells and HEK293 cells expressing various forms of GPR83 (full-length or deletion variant, with or without HA tag) in competition with different concentrations of natural PEN or Tyr-PEN, ranging from 1 pM to 1 μM. Yet, neither of these different assay conditions led to a PEN-GPR83 binding signal above background or a concentration-dependent displacement of the radioligand (Figure 5A-D). In contrast, radiolabeled angiotensin-II ([^125^I]-AII) used as a positive control strongly bound to pancreatic neuroendocrine BON cells expressing the cognate receptor AGTR1 and was displaced by unlabeled AII in a concentration-dependent manner, yielding an IC_50_ of 0.4 nM (Figure 5E).

**Figure 5.**
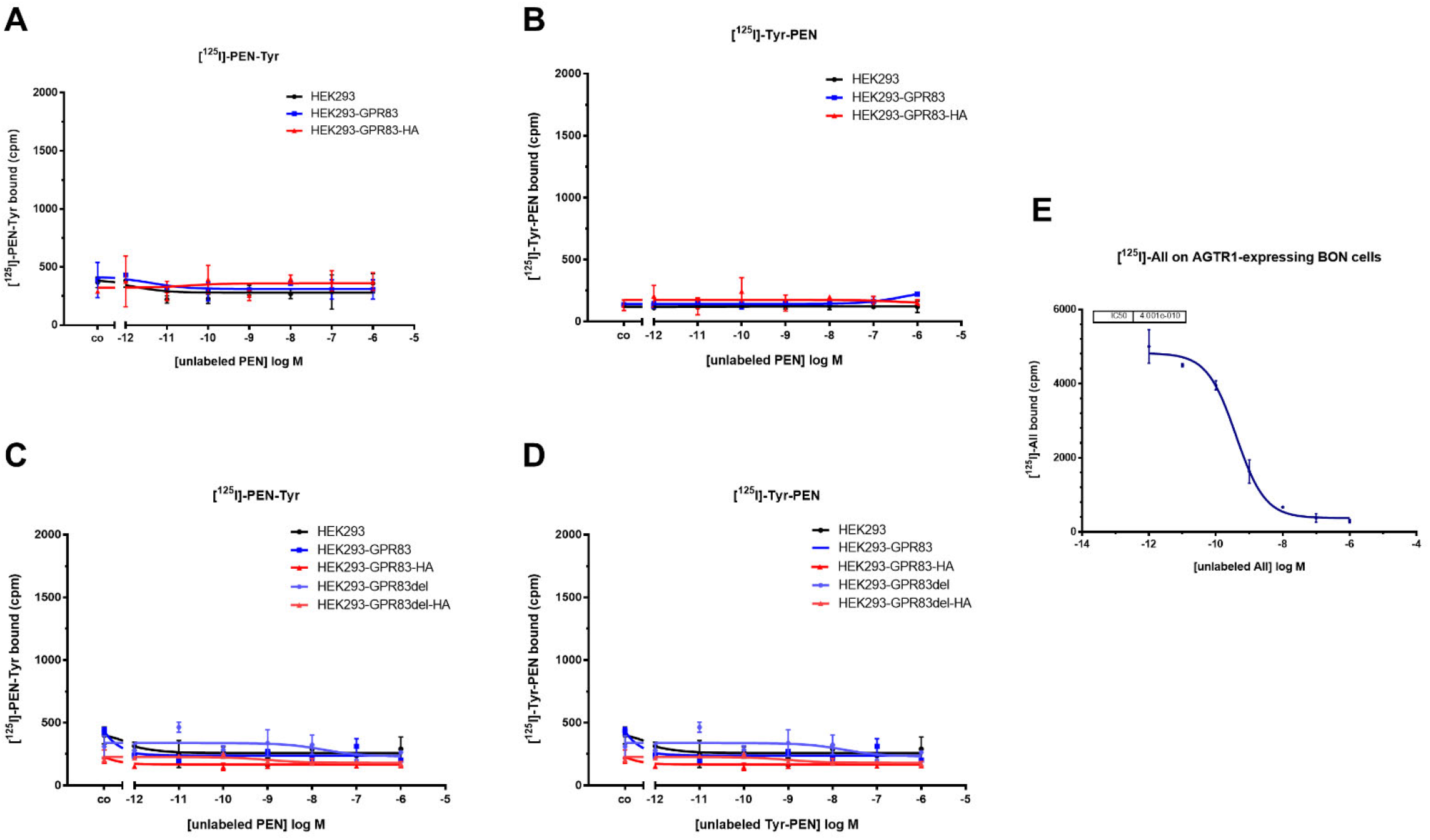
HEK293 wild-type cells and HEK293 cells overexpressing various forms of GPR83 were used in radioligand binding assays with two different radioactively labeled PEN peptides: [^125^I]-Tyr-PEN or [^125^I]-PEN-Tyr. **(A)** Radioligand [^125^I]-PEN-Tyr competed by unlabeled wild-type PEN. **(B)** Radioligand [^125^I]-Tyr-PEN competed by unlabeled wild-type PEN. **(C)** Radioligand [^125^I]-PEN-Tyr competed by unlabeled wild-type PEN. **(D)** Radioligand [^125^I]-Tyr-PEN competed by Tyr-PEN. **(E)** Radioligand binding assay using [^125^I]-AII competed with unlabeled AII on BON cells expressing angiotensin receptor AGTR1

## 3. Discussion

In the present study, evidence for a molecular interaction of the neuropeptide PEN with the cell surface receptor GPR83 could not be found. PEN was unable to specifically activate GPR83 as measured by intracellular calcium mobilization, cAMP production, β-arrestin activation and ERK1/2 phosphorylation assays. While PEN was able to elicit a phosphorylation of tyrosine 202/204 of ERK1/2 as evidenced by Western blotting, the reaction was not specific for GPR83, since wild-type human embryonic kidney HEK293 cells, expressing approximately 100-fold less receptor (Figure 1A) also demonstrated a similar phosphorylation after incubation with PEN (Figure 4A-B). It may therefore be speculated that HEK293 cells endogenously express another receptor for PEN, which is coupled to ERK1/2 activation. A number of relevant controls led to the expected strong increase in intracellular Ca^2+^, yet none of two synthesis batches of PEN from different sources was found to yield a signal above background. This result was confirmed in HEK293 cells as well as in human osteosarcoma cells (U2OS), both transfected with either full-length or truncated GPR83 lacking the N-terminus (Figure 2A-C). Basal signaling by GPR83 reported earlier as evidenced by an IP_3_ increase [19] could also not be confirmed in the Ca^2+^ release assay (Figure 2D). While the initial report on PEN activating GPR83 showed evidence of PEN-induced intracellular Ca^2+^ mobilization in CHO cells transfected with GPR83 and chimeric Gα16/i3 protein [10], this could not be confirmed in this study using either GPR83-transfected HEK293 (± Gα16) or U2OS cells. HEK293 and U2OS cells are of human origin, yet CHO cells are from Chinese hamster (*Cricetulus griseus*) and may confer different coupling option for signaling downstream of human GPR83.

Modification of GPR83-mediated cAMP signaling by PEN has been previously reported [10]. This study, employing multiple clones of GPR83-overexpressing HEK293 and U2OS cells, did not reveal evidence for an increased (via G_s_) or decreased (via G_i_) production of cAMP after incubation with concentrations of up to 1 μM PEN. Forskolin, however demonstrated the expected strong induction of intracellular cAMP (Figure 3). While the earlier study employed CHO cells from *Cricetulus griseus* and Neuro2A cells from *Mus musculus* overexpressing human GPR83 [10], this study used human HEK293 and U2OS cells overexpressing human GPR83 for testing cAMP activity. The different cellular background as well as differences in protocol and detection method may have contributed to contrasting results. As noted in [10], some batches of HEK293 cells had not responded to PEN. Differential evolution of HEK293 lab clones involving signaling components may be relevant here. Another potential mechanism for differences in reactivity would be a differential association of GPR83 with e.g. MRAP proteins, which may give rise to a modulation of GPCR signaling [20,21]. Recently, a pilot screen of a novel peptide hormone library has identified the protein family with sequence similarity 237 member A (FAM237A) as an activator of GPR83-mediated β-arrestin recruitment in murine Att20 and STC-1 cells [14]. PEN was not reported to be tested in this paper. In the current study, the PRESTO-Tango β-arrestin activation assay [18] was employed to test for GRP83 activation by PEN. However, PEN failed to generate a signal in this assay in HTLA-GPR83 cells, while several bradykinin-related peptides yielded strong responses in HTLA-BDKBR2 cells (Figure 4C-D). A common GPCR signaling pathway would involve ERK1/2 activation downstream of β-arrestin recruitment [22,23]. However, the current study could not find indications for a specific GPR83-related activation by PEN of either β-arrestin or ERK1/2.

In addition to the lack of GPR83-mediated signaling, PEN also failed to show any interaction with GPR83 in the radioligand binding assay. Even with a redundant design of the assay on multiple levels (up to four different GPR83-overexpressing cell clones plus two different PEN variants with an added tyrosine on either the amino-or carboxyterminus radiolabeled and used in the assay), no interaction could be detected between PEN and GPR83 (Figure 5). While one end of the peptide may be crucial for binding the receptor, an absolute requirement of both peptide termini to be free of another added amino acid (necessary for radioiodination) seems unlikely. Recently, Li and coworkers have confirmed the interaction of FAM237A, but not of PEN, with GPR83 in multiple luminescence-based interaction and activation assays [16]. Their study, like this one, was unable to provide data for binding or activation of PEN with GPR83. In contrast to the paper by Gomes et al., [10] which used GPR83-transfected hamster CHO and mouse Neuro2A cells for binding assays, the present study employed human HEK293 and U2OS cells transfected with the human receptor. This difference in cellular background may not only give rise to a modulation of downstream signaling via heterodimerization of GPCRs [24-26]. It may also induce a difference in ligand binding, as it may e.g. lead to internalization of a receptor even in the absence of ligand and thus to a loss of binding sites at the cell surface [27]. Heterodimerization has previously been demonstrated for GPR83 and thus might contribute to the observed differences in binding outcome [17]. Furthermore, the use of whole cells versus membrane preparations and other binding protocol differences may have had their influence.

While covering important potential signaling pathways, this study is limited by not including other GPCR-typical signal chains and readouts indicative of a receptor activation. However, the absence of Ca^2+^, cAMP, β-arrestin and ERK1/2 activation, and in particular the lack of binding interaction appears to be notable evidence against PEN engaging with GPR83 in a typical ligand-receptor interaction. Future studies will have to further contribute to bringing in line the conflicting data available so far.

## 4. Materials and Methods

### Reagents and Cell Culture

If not indicated otherwise, chemicals were obtained from Carl Roth (Karlsruhe, Germany) or Sigma Aldrich (St. Louis, MO, USA). GPR83 cDNA clones were subcloned into pcDNA 3.1 from plasmids as published before [17]. An independent GPR83 cDNA clone was taken from the PRESTO-Tango GPCR kit, a gift from Bryan Roth (University of North Carolina, Chapel Hill, NC, USA) (kit #1000000068; addgene, Waterton, USA). BON cells [28] were a kind gift of Courtney Townsend (University of Texas, Galveston, TX, USA). HTLA cells [29] were kindly provided by Gilad Barnea (Brown University, Providence, RI, USA). All other cell lines were purchased from ATCC/LGC Standards (Wesel, Germany). The cell lines were cultured in RPMI 1640 medium (Biochrom AG, Berlin, Germany) or McCoy’s 5A modified medium (Biochrom AG, Berlin, Germany), each supplemented with 10% fetal calf serum (Biochrom AG, Berlin, Germany). All tumor cell lines were cultured in an incubator (Labotect, Rosdorf, Germany) providing a humidified atmosphere at 37 °C with 5% CO_2_. Cells were passaged every 3-5 days.

### Peptide synthesis

Human PEN (designated PEN #1; pro-SAAS 221-242: AADHDVGSELPPEGVLGALLRV) was synthesized on solid support by means of standard Fmoc methodology. Detailed protocol and characterization is provided in the supporting information. A second batch of human PEN (PEN #2) was purchased from Phoenix Pharmaceuticals (Burlingame, USA; cat #004-52). The peptides for radiolabeling (Tyr-PEN and PEN-Tyr) were obtained from peptides&elephants (Hennigsdorf, Germany).

### RNA isolation and reverse transcription quantitative real-time PCR (RT-qPCR)

The preparation of RNA for RT-qPCR experiments was performed according to the manufacturer’s instructions using the RNeasy Plus Mini Kit (Qiagen, Hilden, Germany). Isolated RNA was transcribed into cDNA using the High Capacity cDNA Reverse Transcription Kit (Applied Biosystems, Darmstadt, Germany). For this, 3.2 μg of RNA was transcribed into 80 μL of cDNA (final concentration: 40 ng/μL) according to the manufacturer’s protocol. In addition, as a negative control for each sample, a preparation without reverse transcriptase was analyzed in RT-qPCR. Quantitative real-time human-specific primer sets for CyberGreen PCR amplifications were obtained from TIB Molbiol (Berlin, Germany). Primer sequences were:

hGPR83-Fwd: 5’-TCAAGAACCAGCGAATGCAC-3’

hGPR83-Rev: 5’-TGCTGTTCACAAAGCGAACC-3’

hGAPDH-Fwd: 5’-TGCACCACCAACTGCTTAGC-3’

hGAPDH-Rev: 5’-GGCATGGACTGTGGTCATGAG-3’

hUBC-Fwd: 5’-ATTTGGGTCGCAGTTCTTG-3’

hUBC-Rev: 5’-TGCCTTGACATTCTCGATGGT-3’

For CyberGreen-based PCR, Sso Fast EvaGreen Supermix (Bio-Rad, Düsseldorf, Germany) was used. PCR was performed using the CFX96 real-time system (Bio-Rad). For this purpose, 30 ng of cDNA were added. The analysis was performed using qBase PLUS (Biogazelle, Zwijnaarde, Belgium) according to the ΔΔCt-method. As multiple reference genes were included in the normalization, the geometric mean of the relative housekeeping expression was first determined and then this value was used to normalize the target gene expression. For plotting, data were normalized to a cq value of 34.

### Immunofluorescence

Cells were seeded on glass coverslips and fixed in a 1:1 methanol-acetone mixture for 2 minutes at room temperature, followed by blocking with 5% goat serum in PBS for 30 minutes. Cells were then washed in PBS and placed in a humid chamber to be stained with the primary antibody anti-HA tag at a concentration of 8 μg/mL (anti-rabbit, #H6908, Sigma-Aldrich, Deisenhofen, Germany) in PBS, 0.1 % BSA for 1 h. After washing with PBS, cells were incubated with a Cy3-conjugated goat anti-rabbit secondary antibody (#115165146, Jackson ImmunoResearch, West Grove, PA, USA) at a concentration of 1.5 μg/mL in PBS 0.1 % BSA for 30 min at room temperature. After washing with PBS, and staining with DAPI (200 nM in PBS, 5 min, followed by a PBS wash), coverslips were briefly dipped into 96 % ethanol, air-dried and mounted on glass slides with ImmuMount (Thermo Fisher Scientific, Waltham, MA, USA). Mounted cells were imaged using an inverted microscope (CellObserver Z1, Carl Zeiss, Jena, Germany) with a 63x NeoFluar oil immersion objective.

### Calcium mobilization assay

Cells were seeded at 50,000 cells/well in a poly-d-lysine coated 96-well black well/clear bottom plate (BD Falcon, Franklin Lakes, USA) and cultured overnight. 16-24 hours later, cells were loaded with Fluo4-AM (Invitrogen, Waltham, USA) for 30 min. Cells were then washed two times with assay buffer C1. After the final wash, a 100 μL residual volume remained on the cells. Peptides and other ligands were dissolved in 10% DMSO to a concentration of 1 mM and were diluted in C1 solution with 0.1 % BSA. They were aliquoted as 2x solutions in 96-well plates and were transferred simultaneously by the robotic system within the imager from the ligand plate to the cell plate. Fluorescence was recorded with the fluorescence imaging plate reader CellLux (Perkin Elmer, Waltham, USA) simultaneously in all wells at an excitation wavelength of 488 nm and emission wavelength of 510 nm in 1.5 s intervals over a period of 4 min. Fluorescence data were generated in duplicate and experiments were repeated at least two times.

### cAMP assay

To measure intracellular cAMP production, the LANCE Ultra cAMP kit (Perkin Elmer) was used according to the manufacturer’s instructions. The day before experiment, cells were seeded onto a 96-well plate with approximately 80% confluence. After 16-24 hours, medium was removed and 100 μL/well of serum-free medium was added, followed by a 3-hour incubation. Thereafter, medium was removed and 40 μL of ligand dissolved in cAMP stimulation buffer was pipetted to cells, then incubated for 10 min. The ligands used were pipetted in duplicate. For concentration-response curves, serial dilutions were carried out from the stock solutions of 1 mM to create a concentration range from 1 μM to 30 pM. As controls, 4 wells were treated with buffer only (negative control) and 4 wells with 50 μM forskolin (positive control). After a 10 min incubation, 10 μL of a 0.5% Triton-X 100 solution in PBS are added to each well. To lyse cells, the plate was shaken on a Vibramax 100 Shaker (750 rpm) at room temperature for 10 min. 10 μL of the cell lysates were transferred into a white 384 well plate (ProxiPlate, PerkinElmer). 5 μL of the Europium-cAMP tracer (1 to 50 dilution of Eu-tracer stock in cAMP detection buffer) was added to each well. Next, 5μL of the ULight-anti-cAMP solution (1 to 150 dilution of ULight-tracer stock in cAMP detection buffer) added to each well. After a 1 h incubation period at room temperature in the dark the TR-FRET signal was measured using an EnVision 2103 Multilabel Reader (Perkin Elmer).

### ERK1/2 phosphorylation assay

Effects of GPCR ligands on MAPK signaling were assessed by western blot analysis of lysates from stimulated cells. The ERK1/2 activation assay was performed as described before [30]. Briefly, 50,000 cells per well were seeded onto adherent 6-well plates and cultured until cells were 80 % confluent. After incubation with ligands (10 μM PEN #1, 100 ng/mL EGF or 200 nM PMA in 0.1 % FCS/DMEM) for 15 min, cells were lysed using 200 μL/well cell lysis buffer (100 mM Tris-HCl pH 8.8, 1 % SDS) and protease inhibitors (cOmplete, Roche) on ice for one h. Immunolabeling was realized using the rabbit monoclonal primary antibody anti-phospho-ERK_1/2_ Thr^202^/Tyr^204^ (#4370, Cell Signaling Technologies, 1:2,000), or anti-total ERK_1/2_ (#05-1152; millipore, Burlington, USA; 1:2,000), respectively. Blots were analyzed using densitometric quantification using Image Lab Software v4.1 (Bio-Rad).

### TANGO assay

For the determination of β-arrestin recruitment after GPCR activation, the PRESTO-Tango system was utilized [18]. On day one, HTLA cells (HEK293 cell line stably expressing a tTA-dependent luciferase reporter and a β-arrestin2-TEV fusion gene, maintained in RPMI supplemented with 10% FBS, 2 μg/mL puromycin and 100 μg/mL hygromycin) were seeded into 150 mm cell culture plates at 10 x 10^6^ cells per well in 30 mL of medium. The next day, transfections were performed directly in the 150 mm plates using jetPEI (Polyplus, Illkirch, France). Plasmid DNA of interest was prepared for transfection following the jetPEI protocol and with a DNA:jetPEI ratio of 1:1. A GFP-encoding plasmid was added as a transfection control (GFP plasmid added at 10% of total DNA to the total NaCl used for plasmid DNA dilution). On day 3, transfected cells were transferred at 15,000 cells per well in 30 μL of medium into 384-well black well, clear bottom plates. On day 4, ligand library multiwall plates (10 μM) were thawed and 23 μL were added to each well. On day 5, 50 μL per well of BrightGlo luminescence reagent (diluted 10-fold with HBSS buffer) were carefully added and plates were incubated for 10 min before luminescence reading in the EnVision 2103 Multilabel Reader (Perkin Elmer).

### Radioiodination

Two variants of human PEN were utilized for radioiodination in this study: one with a tyrosine added at the aminoterminal end (Tyr-PEN: YAADHDVGSELPPEGVLGALLRV-NH_2_), and one with an additional tyrosine attached to the carboxyterminal end (PEN-Tyr: AADHDVGSELPPEGVLGALLRVY-NH_2_). Both were labelled with ^125^I using the chloramine-T method. Chloramine-T and sodium metabisulfite solutions were freshly prepared in water. For labelling, 10 μL of peptide (stock 1 mM) with 15 μL of sodium phosphate buffer (0.5 M, pH 7.6) was mixed with 37 MBq carrier-free Na^125^I (NEZ033L010MC, Perkin Elmer, Waltham, US). 4 μl chloramine T (1 mg/mL) were added to start the reaction and after 30 seconds, 4 μl sodium metabisulfite (2 mg/mL) were added to stop the iodination. Labelled radioactive peptide was separated from unlabeled peptide by HPLC purification (Agilent ZORBAX 300 Extend-C18 column) using a gradient from 20-50% acetonitrile (+0.1 % TFA) against water (+0.1 % TFA) for 20 min. To determine the retention time of the radioactive peptide, 1-2 μL of the reaction mixture were analyzed before the purification run. The fraction containing the radiolabeled peptide peak was then collected, diluted with binding buffer to prevent radioautolysis, aliquoted and stored at -80 °C.

### Radioligand binding assay

For competitive radioligand binding assay, approximately 50,000 cells/well were seeded in a 96 well plate incubated overnight at 37°C. Next day, cells were incubated in binding buffer (50 mM HEPES pH 7.4, 5 mM MgCl_2_, 1 mM CaCl_2_, 0.5 % BSA, cOmplete protease inhibitors) containing 100,000 counts per minute ^125^I-labelled peptide and increasing concentrations of unlabeled peptide. After 2 h of incubation, cells were washed for 2-3 times with ice-cold washing buffer (50 mM Tris-HCl, pH 7.4, 125 mM NaCl, 0.05% BSA) and lysed with 1 N NaOH (80 μL/well). The lysed cells were transferred to vials and measured in a gamma counter (Wallac 1470 Wizard, Perkin Elmer, Waltham, MA, USA). Obtained cpm values were plotted with GraphPad Prism 7 and the data were fitted using nonlinear regression (one site - fit Ki, one site - fit logIC_50_).

## Supporting information

Synthesis information

## Author Contributions

Conceptualization, G.K., B.K., C.G.; methodology, Y.G., V.A., V.S., C.G.; formal analysis, Y.G., V.A., V.S., C.G.; investigation, Y.G., V.A., V.S., C.G.; resources, B.K., P.S.; writing—original draft preparation, C.G.; writing—review and editing, V.A., V.S., G.K., P.S., C.G.; supervision, B.K., P.S., C.G.; funding acquisition, P.S., C.G. All authors have read and agreed to the published version of the manuscript.

## Data Availability Statement

The quantitative data for this study have been permanently published in a public repository accessible via this link: https://doi.org/10.5281/zenodo.8305579

