## Supplementary material for "Is the neuropeptide PEN a ligand of GPR83?": Synthesis information

Applied Materials

Fmoc-L-amino acids were purchased from ORPEGEN Peptide Chemicals GmbH (Heidelberg, Germany). Fmoc-Val-Wang resin was from Novabiochem (Merck Chemicals GmbH, Darmstadt, Germany). All solvents were used from VWR (Darmstadt, Germany) without further purification. All other chemicals were bought from Acros (Geel, Belgium), VWR (Darmstadt, Germany) or Merck (Darmstadt, Germany) at highest commercially available purity and used as such.

Peptide synthesis and purification:

Peptides were synthesized with an Activo-P11 automated peptide synthesizer (Activotec, Cambridge,United Kingdom) in a 0.05 mmol scale on a solid support by standard Fmoc methodology on a preloaded Fmoc-Val-Wang resin (0.32 mmol/g loading) using a 10 mL polypropylene reactor at rt. Fmoc-deprotection was performed with 20 % piperidine + 0.1 M 1-hydroxybenzotriazole (HOBt) in DMF (3 x 10 min). The amino acids were introduced in a double coupling with five equivalents excess with respect to the amount of resin using 4.9 eq. of HATU as coupling reagent and 10 eq. DIPEA for 30 min. In order to avoid aspartimid-formation Fmoc-Asp(OEpe)-OH (Merck Chemicals GmbH, Darmstadt, Germany) was used in all Asp positions.

After synthesis the peptide were cleaved from the resin by treatment with trifluoroacetic acid (TFA) containing triisopropylsilane (TIS,5 % (v/v)), water (5 % (v/v)) [1 mL cleavage-cocktailper 50 mg resin] for 2.5 h at rt. Afterwards the resin was washed twice with TFA (2 mL) and DCM (2 mL) and excess solvent was removed by evaporation. The crude peptide was then precipitated with ice-cold diethyl ether (2 x 40 mL), and after centrifugation dried by lyophilization. The obtained crude peptide was purified on a LaPrepΣ low-pressure HPLC system (VWR, Darmstadt, Germany) using a Kinetex RP-C18 endcapped HPLC column (5 µM, 100 Å, 250 × 21.2 mm, Phenomenex®, USA). A Security GuardTM PREP Cartridge Holder Kit (21.20 mm, ID, Phenomenex®, USA) served as pre-column. As eluents deionized water (Milli-Q Advantage® A10 Ultrapure Water Purification System, Millipore®, Billerica, MA, USA) and acetonitrile (ACN), both containing 0.1% (v/v) TFA were applied. HPLC runs were performed with a flow rate of 15.0 mL/min, UV detection occurred at 220 nm. A linear gradient of 5–100 % ACN in water (both + 0.1 % TFA) was applied within18 min. Data analysis was conducted with an EZChrom Elite-Software (Version 3.3.2 SP2, Agilent Technologies, Santa Clara, CA, USA). The fractions containing pure peptide were combined, and ACN was removed by rotary evaporation. Lyophilization of the remaining aqueous solution yielded 20 mg of the pure peptide as a white powder.

Peptide characterization

The purity of the peptide was controlled by analytical HPLC on a Chromaster 600 bar DAD-System with CSM software (VWR/Hitachi, Darmstadt, Germany) with a Kinetex C18 column (5 µm,250 Å~ 4.6 mm, Phenomenex®, Torrance, CA, USA) with a linear gradient of 5-70 % ACN + 0.1 TFA in 18 min. The Chromaster system works with a low-pressure gradient containing a HPLC-pump (5160) with a 6-channel solvent degaser, an organizer, an autosampler (5260) with a 100 µL sample loop, a column oven (5310) and a diode array flow detector (5430). High resolution mass spectra were recorded on an Agilent 6220 ESI–ToF LC–MS spectrometer (Agilent Technologies Inc., Santa Clara, CA, USA) to identify the pure peptide. The sample was dissolved in a 1:1 mixture of water and ACN containing 0.1% (v/v) TFA and injected directly into the spray chamber by a syringe pump using a flow rate of 15 µL min$^{-1}$. A spray voltage of 3.5 kV was used; the drying gas glow rate was set to 5 L min$^{-1}$ and the nebulizer to 30 psi. The gas temperature was 300 °C. Peptide mass calculator v3.2 (http://rna.rega.kuleuven.be/masspec/pepcalc.htm) was applied to obtain the calculated (calc.) monoisotopic values for the desired peptide.

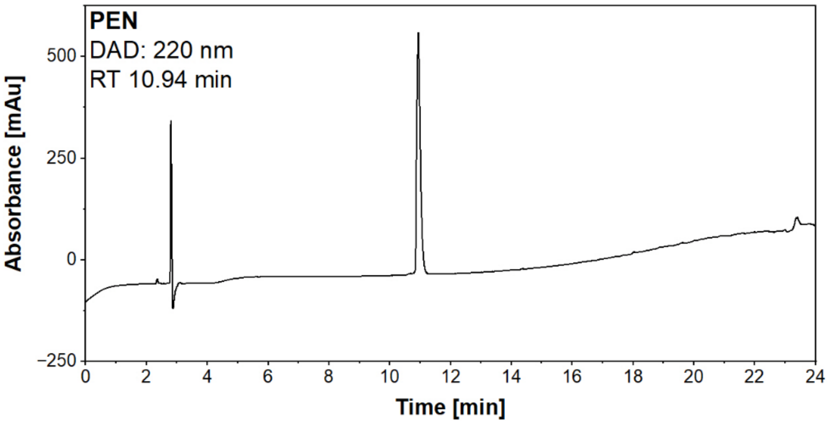


*Figure S1: Analytical HPLC chromatograms of purified peptides; column: Kinetex C18 column (5 µm,250 Å~ 4.6 mm, Phenomenex®, Torrance, CA, USA; Solvent A was $H_2O$, solvent B was acetonitrile, both containing 0.1% (v/v) TFA. The flow rate was 1 mL/min; linear gradient from 5% B to 70% B over 18 min. The desired compound eluted at a retention time (RT) of 10.94 min. Identification by ESI–ToF mass spectrometry: $[M\text{-}1H]^{1+}$ 2215.1721 (calc.) $[M\text{-}1H]^{1+}$ 2215.1384 (obs.), $[M\text{-}2H]^{2+}$ 1108.0900 (calc.) $[M\text{-}1H]^{1+}$ 1108.0985.1384 (obs.).*
